# On the regulation of photosynthesis in pea leaves exposed to oscillating light

**DOI:** 10.1101/2022.03.01.482448

**Authors:** Dušan Lazár, Yuxi Niu, Ladislav Nedbal

## Abstract

Plants grow in nature often in fluctuating irradiance. In the laboratory, the dynamics of photosynthesis is usually explored by instantaneously exposing dark-adapted plants to constant light and detecting the dark-to-light transient, which is only a poor approximation of natural phenomena. Aiming at a better approximation, we exposed pea leaves to oscillating light and measured, during oscillations, changes in function of photosystem I and II and of the proton-motive force at the thylakoid membrane. The dynamics depends on the oscillation period, leaving information about the regulatory networks. As demonstrated for selected period of the oscillation of 60 s, the regulations try to keep reactions centres of photosystems I and II open. A possible evaluation of obtained data is presented and involvement of particular processes in regulation of photosynthesis is discussed. The forced oscillations provide information-rich fingerprint of complex regulatory networks. Further progress in understanding the networks is expected from experiments involving chemical interventions, plant mutants, and by using mathematical modelling and the system identification and system control tools, as already applied in other parts of science.

**Highlight:** Measurement of photosynthetic signals during illumination of plants by light, whose intensity oscillates as sinus function provides information about regulation of photosynthesis in fluctuating light.

## Introduction

Adaptation of the photosynthetic apparatus to contrasting and, often, extreme environmental conditions, including dynamically changing light intensity (e.g., Yamori, 2016) has been one of the important diversifying factors in the evolution of plants. To deal with the changing environment, plants developed elaborate and diverse regulations (e.g., Tikhonov, 2015) to optimize the yields and minimize the potential damage by harmful by-products (e.g., Pospíšil, 2016). Increasing our understanding of the dynamics of photosynthetic regulation in nature requires techniques that can investigate plants in fluctuating light (e.g., Rascher and Nedbal, 2006; Matsubara, 2018; Gjindali et al., 2021). Simulations of the natural random light fluctuations and approximations, e.g., by repeated step changes in light intensity are appearing in recent literature with a growing frequency (e.g., Cruz et al., 2016; Annunziata et al., 2017; Vialet-Chabrand et al., 2017; Adachi et al., 2019; Li et al., 2019).

An alternative approach using regular, harmonically oscillating light was introduced by Nedbal and Březina (2002) in analogy to the frequency-domain analyses that are widely used in physics and engineering. Any fluctuating light, random or regular, can be represented by a weighted sum of elemental harmonic components of characteristic frequencies (Schwartz, 2008) and one may, thus, reduce the complexity of such an investigation to exposing plants to light that is sinusoidally oscillating with a single frequency, called harmonic modulation. The angular frequency of the modulation 2π/T, where T is period of the oscillating light, can be gradually changed to cover the whole range of characteristic dynamic components as they occur in a particular natural environment. Such a frequency-domain analysis has been already applied in plant research (e.g., Nedbal et al., 2003; 2005) and, recently, supported by mathematical models and further experiments (Nedbal and Lazár, 2021). Since course of measured photosynthetic signal forced by the sinusoidal illumination deviates from the course of illumination, response of the measured signal to illumination is thus non-linear. The nonlinearity can be constitutive, i.e. directly related to primary functions of explored system, and regulatory nonlinearity (Bich et al., 2016). The regulatory nonlinearity manifest itself as presence of the so-called upper harmonics (multiplies of the basal frequency of modulation, i.e., 2*2π/T, 3*2π/T, etc.) in the measured signal (Nedbal and Lazár, 2021), i.e., by other words, if the measured signal is not saturated, additional waves and/or bumps appear in the signal. The harmonically oscillating illumination, thus, yields not only insights into elemental dynamics of photosynthesis in fluctuating light but also reveals frequency domains that characterize various mechanisms of photosynthesis regulation (Nedbal and Březina, 2002; Nedbal et al., 2003; 2005; Nedbal and Lazár, 2021).

The response of plants to changing light is often sensed by measuring the chlorophyll (Chl) fluorescence (ChlF) (reviewed in Bąba et al., 2019). A main part of variable ChlF in the initial rise of the ChlF induction that occurs with the exposure of dark-acclimated plants to constant light is attributed to the progressing photochemical reduction of the first quinone electron acceptor, Q_A_, of photosystem II (PSII) (reviewed in Lazár, 1999; Stirbet and Govindjee, 2012). However, even Q_A_ is already reduced, i.e., the reaction centres (RC) of PSII (RCII) are closed, the ChlF can further increase and the increase is probably caused by light-induced conformational changes of PSII and/or the electric fields (Vredenberg and Bulychev, 2002; Magyar et al., 2018; Laisk and Oja, 2020; Sipka et al., 2021). The latter rise of ChlF accounts for about 1/3 of ChlF measured with saturating light.

The photochemical alteration of ChlF occurs also in harmonically modulated light, in which some of the PSII RCs (RCII) may open around the light minima and close around the light maxima. The respective fractions of open and closed RCIIs are expected to change with the frequency and amplitude of the harmonic light modulation, providing precious information about the dynamic of the primary processes and regulation in fluctuating light.

Later phases of the ChlF induction are typically attributed to the activation of the CO_2_ assimilation in the Calvin-Benson cycle and to non-photochemical quenching (npq) of ChlF. The npq is a generic term that, strictly speaking, includes also processes that decrease the photosynthetic energy conversion and ChlF emission in a strong light by lowering the effective absorption cross-section of the RCIIs rather than by an excitation quenching by heat dissipation. More typically, however, npq is attributed to the dissipation of the excess excitation energy as heat (Papageorgiou and Govindjee, 2014). In plants, this quickly reversible, ‘high-energy’ npq (qE) is related to acidification of lumen in light, formation of carotenoid zeaxanthin and antheraxanthin from violaxanthin, and requires presence of PSII protein subunit S (reviewed in Holzwarth and Jahns, 2014).

One may presume that the Calvin-Benson cycle remains active in the harmonically oscillating light as long as the light minima are not too long and not too close to darkness. On the other hand, the npq-regulation is likely to modulate the ChlF emission response to an extent determined by the interference between light modulation frequency and amplitude on one hand, and by the activation and deactivation characteristics of the npq processes. We presume that this interference represents a unique opportunity to dynamically separate and identify individual npq mechanisms.

Another tool to dissect various contributions to ChlF modulation are multiple-turnover saturating pulses (MTSP) that can transiently close the RCIIs by transiently congesting the electron transport pathways. The MTSPs are routinely applied in the time-domain measurements with constant light (reviewed in Lazár, 2015). However, by applying the MTSPs, one has to be aware that the maximal ChlF determined by means of MTSPs might reflect to some extent also the effect(s) not related to the photochemical closure of the RCIIs (see above).

The information obtained with ChlF can be complemented by the dynamics of transmittance optical proxy I830. The I830 signal, measured as a difference of the leaf transmittance at 875 and 830 nm, reflects mainly the redox dynamics of the primary electron donor P700 in photosystem I (PSI) (e.g., Klughammer and Schreiber 1991) although redox changes of plastocyanin and ferredoxin may also contribute. Similarly to the ChlF, additional information about PSI function can be obtained by application of MTSPs (e.g., Schreiber and Klughammer, 2008a). The Dual-PAM instrument (Heinz Walz GmbH, Effeltrich, Germany), used in the present study, can measure ChlF and I830 signal simultaneously. The instrument can provide yet another important complementary insight into photosynthetic dynamics by measuring the difference of the leaf transmittance at 550 and 515 nm, named P515 (e.g., Schreiber and Klughammer, 2008b). This signal reflects the electrochromic shift of carotenoids and Chl b absorption bands that is caused by the electric potential difference, ΔΨ, across the thylakoid membrane (TM). A contribution of other effects to the P515 signal, i.e., absorption change at 535 nm (see, Ruban et al., 2002) and at 550 nm (see, Van Wittenberghe et al., 2019) is eliminated by calculating the transmittance difference at the two wavelengths, and can be distinguished as a change in baseline in time scale different than changes in P515, respectively (Schreiber and Klughammer, 2008b). Enhancing further the information, the relaxation of the P515 signal that occurs when the actinic light is switched off can be used to separate and quantify the chemical (ΔpH) and electric (ΔΨ) components of the proton motive force (pmf) over the TM (Cruz et al., 2001; Schreiber and Klughammer, 2008b).

The information content of ChlF, and I830 reporter signals can be further enriched by applying MTSPs at different phase-points of the light oscillation (Nedbal et al., 2003), in the same way as it is routinely done during the induction in the dark-to-light transition mentioned above. Similarly, the P515 relaxation in different phases of the light oscillation can be also recorded by abruptly switching the actinic light off.

In this work we provide detailed information about measurement of the forced oscillations in ChlF, I830, and P515 signals, and of procedure for measurement of the quantum yields and other parameters during the forced oscillation of the signals. We show the forced oscillations of the signals of pea leaves caused by illumination of the leaves by red actinic light oscillating with periods ranging from 1 s to 300 s. For the period of 60 s, we show quantum yields related to the function of PSII and PSI as well as changes of pmf and its ΔpH-dependent and ΔΨ parts changing with the light oscillation, the latter presented for the first time. Further, we show a possible analysis of some of the parameters. By comparing the evaluated parameters, we infer on the mechanisms of the regulation of photosynthetic function in fluctuating light. To support our conclusions, more work with different mutants and/or chemical interventions (inhibitors, electron acceptors and donors, etc.) should be done in future to better understand mechanisms of photosynthesis regulation in fluctuating light.

## Material and methods

### Plant material

Seeds of pea (*Pisum sativum*) were sown to pots with perlite and Knop solution and placed in a growth chamber (Percival AR-100L3, Percival Scientific, USA) under controlled conditions of 16 h white light (150 μmol (PAR photons) m^−2^ s^−1^) and 8 h dark. The temperature was kept between 22 and 20 °C and the relative air humidity was 60% during the germination and growth. Pots with 15 to 20 days old pea plants were removed from the growth chamber and kept in darkness for 30 minutes before the measurements. Well-developed green leaves attached to the plants were, one by one, gently inserted between the optical heads of the instrument and measured.

### Methods

#### The used instrument, light colours and intensities

The Dual-PAM-100 (Heinz Walz GmbH, Effeltrich, Germany) instrument with the optical heads DUAL-E and DUAL-DB was used for simultaneous measurements of the ChlF and I830 on the same leaf. The P515 reporter signal was consecutively measured on another leaf using the DUAL-EP515 and DUAL-DP515 optical heads.

Red (635 nm) actinic light was generated by the same instrument with its intensity sinusoidally oscillating between 0 and 250 μmol (photons) m^−2^ s^−1^. This intensity range was chosen as leading to the most pronounced non-linear upper harmonic modulation (i.e., the measured output signal is described by a sum of sinus functions whose periods, in addition to T, are T/2, T/3, …, where T is period of input sinusoidal light) that is a hallmark of regulatory non-linearity (Nedbal and Březina, 2002; Bich et al., 2016; Nedbal and Lazár, 2021). ChlF was excited by blue measuring flashes (460 nm) and detected by the pulse amplitude modulation (PAM) method. I830 and P515 were detected as the differences of the leaf transmittances at 875 and 830 nm, and at 550 and 515 nm, respectively. MTSPs of white light (intensity of 5000 μmol (PAR photons) m^−2^ s^−1^, duration of 0.3 s) and illumination by far-red light (720 nm, duration of 9 s) were also generated by the optical heads of the same instrument. For the purpose of a theoretical evaluation, the intensity of oscillating light is converted to values of the rate constant of closure of the reaction centres of photosystem I and II, kL, assuming 1 : 1 ratio for the intensity of excitation light in µmol photons m^-2^ s^-1^ to value of the rate constant in s^-1^. This is based on previous calculations (Lazár and Pospíšil, 1999). Thus, kL was changing from 0 to 250 s^-1^.

#### Measurement of the oscillation itself

The experimental protocol for measurement the oscillation itself started always by exposing a pea leaf for 5 s to measuring flashes to obtain optical signals in the dark-acclimated state of the plant. Subsequently, the leaf was illuminated by the actinic red light of a constant intensity, 250 μmol (photons) m^−2^ s^−1^, for 10 s so that photosynthetic induction starts under constant light illumination. After that, the light oscillations were initiated, first, with irradiance decreasing from the maximum (250 μmol (photons) m^−2^ s^−1^) to a minimum (0 μmol (photons) m^−2^ s^−1^) along a cosine function. The first 60 oscillations, each 1 second long, were performed in the first minute, followed by three minutes, in which another 60 oscillations, each three seconds long, were given. Without interruption, more sets of light oscillations were continued, always with only five periods, T = 10, 20, 30, 60, 120, 180, and 300 s each. This entire sensing protocol was repeated 3 – 4 times, always using a new dark-acclimated plant. Transients shown in Fig. 1 were obtained by averaging the signals in the last two oscillation periods, in which the signals followed sustained, reproducible dynamic patterns ChlF(t/T) and I830(t/T), and P515 (t/T); for an example, see Fig S1 in the supplementary material.

**Figure 1.**
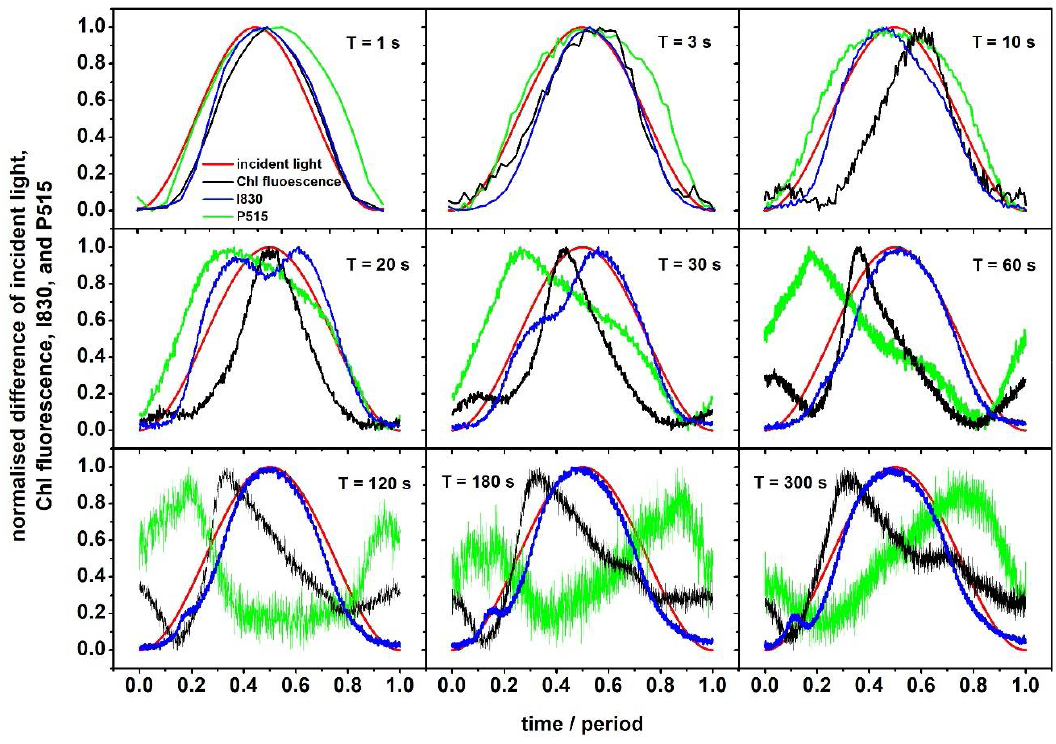
Dynamics of normalized difference of chlorophyll (Chl) fluorescence, I830 and P515. The pea leaves were exposed to oscillating red light with maximal intensity of 250 µmol of photons m^-2^ s^-1^ (also shown) and different periods (T). For details, see Material and methods.

An example of the measured signals is shown in the supplementary material in Fig. S1. The figure shows what the amplitude of the forced oscillations was compared to the initial magnitude of the signals caused by the illumination by the constant light intensity. For the case of P515, a drift of the signal is present. This drift probably reflects changes of zeaxanthin absorption (with maximum at 550 nm) caused by light-induced structural transition of light harvesting complexes of PSII where zeaxanthin was already bound; these changes are not connected with changes in the npq (Van Wittenberghe et al., 2019). We note that the drift in P515, but also in I820 signal, was present occasionally and with a different extent.

#### Measurement for evaluation of the quantum yields

For determining the quantum yields, a leaf from another plant was first illuminated only by the measuring flashes, and minimal ChlF (F_0_) and minimal I830 (P_0_) for the dark-acclimated state were measured. A MTSP was given to transiently reduce the acceptor side of PSII and determine the maximal ChlF (F_M_) characterizing the dark-acclimated state. Following a short darkness, far-red light was used to oxidize the electron transport chain on the donor side of PSI, followed by another MTSP to oxidize fully its P700 primary donor and determine the maximal I830 (P_M_). These are standard routines for determination of the F_0_, F_M_, P_0_, and P_M_ values.

After determining the above mentioned extreme values in dark-acclimated plants, the actinic red light of the constant intensity of 250 μmol (photons) m^−2^ s^−1^ was switched on for 10 s to induce the light acclimation. The oscillations in actinic light intensity then started down from the maximal value, approximating the cosine function. Four-and-half oscillations each with a period of 60 seconds induced a largely stationary dynamic pattern, and measurement of quantum yield was then initiated by giving the first of the MTSPs at the beginning of the oscillation. The reporter signals just before the MTSP were denoted Ft(0s/60s) and Pt(0s/60s), respectively, and the signals read at the end of the MTSP F_M_’(0s/60s) and P_M_’(0s/60s). The second MTSP was given 63 s later, in the next oscillation, phase-delayed by 3 s from the preceding one, and yielding thus signals Ft(3s/60s), F_M_’(3s/60s) and Pt(3s/60s), P_M_’(3s/60s). By repeating this measuring algorithm in the next 19 oscillations, we obtained the signals evenly covering every 3 s the whole oscillation period, ending with Ft(60s/60s), F_M_’(60s/60s) and Pt(60s/60s), P_M_’(60s/60s) from the last MTSP in the last period. Having only one MTSP per each of the 21 one-minute oscillation periods ensured that the MTSPs did not have significant impact on the measured dynamics. This was also proved by always the same shape of the oscillation during the 21 periods. An example of the measurement is shown in the supplementary material in Fig. S2. This quantum yield measurement was repeated with leaves of four plants to obtain mean values and standard deviations of the measured parameters (Fig. 3).

#### Measurement for evaluation of partitioning of the pmf

Similarly, the dynamic of the P515 signal was complemented by separate experiments in which the actinic light (intensity range same as above) was switched off at different points (phase shifts) along the actinic light oscillation period. The protocol always consisted first of nine-and-half oscillations, each 60 s long to induce a stationary P515 dynamic pattern and, during following periods, the light was turned off for 60 s at various phases of the oscillation. The P515 value just before the actinic light was turned off was Et. In dark, the P515 signal first sharply decreased to its minimum, E_min_, followed by a slower increase to a maximum, E_max_. The actinic light was subsequently switching off with different phase-delays appearing in 6 s steps, so that datasets {Et(0/60s), E_min_(0/60s), E_max_(0/60s)}, {Et(6/60s), E_min_(6/60s), E_max_(6/60s)}, {Et(12/60s), E_min_(12/60s), E_max_(12/60s)}, …, {Et(60/60s), E_min_(60/60s), E_max_(60/60s)} were obtained. Before each phase-delayed dark interval, the sinusoidal actinic light was on (from its maximal value) for 2 and half periods, which was enough to achieve the same stationary periodical changes of P515 signal again. An example of the measurement is shown in the supplementary material in Fig. S3. This assay was repeated with leaves from three different plants to determine mean values and standard deviations of the respective parameters (Fig. 3).

#### Evaluated parameters

The primary data on ChlF and I830 obtained by the protocols described above were used to calculate characteristic quantities (Klughammer and Schreiber, 1994; Hendrickson et al. 2004; Schreiber and Klughammer, 2008a; Lazár, 2015) that can be used for a direct molecular interpretation. The quantities are defined as follows:

- the effective quantum yield of PSI photochemistry Y(I) = (P_M_’ - Pt)/(P_M_ - P_0_)
- the quantum yield of non-photochemical quenching of PSI excitation energy due to a limitation on the PSI donor side, Y(ND) = (Pt - P_0_)/(P_M_ - P_0_)
- the quantum yield of non-photochemical quenching of PSI excitation energy due to a limitation on the PSI acceptor side, Y(NA) = (P_M_ - P_M_’)/(P_M_ - P_0_)
- the effective quantum yield of PSII photochemistry, Y(II) = (F_M_’ - Ft)/F_M_’
- the quantum yield of constitutive non-regulatory non-photochemical quenching of PSII excitation energy, Y(f,D) = Ft/F_M_
- the quantum yield of light-induced regulatory non-photochemical quenching of PSII excitation energy, Y(NPQ) = (Ft/F_M_’) - (Ft/F_M_)
- the non-photochemical quenching parameter, NPQ = (F_M_ - F_M_’)/F_M_’ = Y(NPQ)/Y(f,D)?Some of the parameters above were arithmetically mutually dependent: Y(I) + Y(ND) + Y(NA) = 1 and Y(II) + Y(NPQ) + Y(f,D) = 1.

The coefficient of photochemical quenching of PSII excitation energy, assuming energetically connected PSII units, qCU, the minimal ChlF for the light-acclimated state, F_0_’, and the effective quantum yield of alternative electron transport (AET), Y(AET), were calculated from the measured values in Microsoft Excel according to Kramer et al. (2004), Exborough and Baker (1997), and Yamori et al. (2011), respectively, as follows:

- qCU = (F_M_’ - Ft)/{[(p/(p - 1)](Ft - F_0_’) + F_M_’ - F_0_’}, where p = 0.55 (Joliot and Joliot, 1964) is the probability of energy transfer between the connected PSII units
- F_0_’ = F_0_/(((F_M_ - F_0_)/F_M_) + (F_0_/F_M_’))
- Y(AET) = Y(I) - Y(II).

The primary data on the P515 signal were used for estimating changes of pmf and its partitioning into its chemical (ΔpH-dependent) and electrical (ΔΨ) components. By modifying the approach developed for continuous constant light and a steady-state (Cruz et al., 2001; Schreiber and Klughammer, 2008b) we used for the pmf and its partitioning in oscillating light the following formulae:

- pmf = Et – E_min_
- ΔpH-dependent part of pmf = E_max_ – E_min_
- ΔΨ = Et – E_max_

This approach enables evaluation of changes (in relative units) of the pmf itself and its parts (not as fractions of pmf) during the course of oscillating actinic light intensity.

## Results

The dynamic patterns of ChlF, I830, and P515 signals shown in Fig. 1 were detected in pea leaves exposed to red actinic light that was oscillating between 0 and 250 μmol (photons) m^−2^ s^−1^ with periods T ranging from 1 s to 300 s. The dynamics in Fig. 1 represent stationary patterns, i.e., the shape of the oscillations is sustained over longer periods under the oscillation patterns, which were achieved after acclimation to several periods of light oscillation with the respective period T (see Material and methods). Since the sinusoidal oscillating light forced the measured signals to oscillate, we further refer to measured signal changes as forced oscillations. This is in agreement with standard textbooks (Stanford and Tanner, 1985) as well as with papers describing the phenomenon in the plant research (Nedbal and Březina, 2002; Nedbal et al., 2003; 2005; Nedbal and Lazár, 2021).

For the two shortest periods of 1 and 3 seconds, the signals followed closely the light intensity that was forcing the processes (red line). ChlF (black line) and I830 (blue line) responded to the increasing light in the first half period with a slight delay while the rise of P515 (green line) was nearly simultaneous to the light. In contrast, the decline of P515 in the second half period followed the declining light with a significant delay, indicating that a deviation of from course of oscillating light in this frequency domain is stronger for P515 than for the reporters on the primary reactions in PSII (ChlF) and PSI (I830). Activation of mechanisms leading to the P515 rise was nearly instantaneous while deactivation occurred with a delay.

The P515 became convoluted with the periods increasing between 20 to 60 s when the signal was reaching its maximum when the light was still increasing. This trend lead to a further differentiation with periods further increasing from T = 120 to 300 s, resulting in a broad signal depression when the light intensity was high.

ChlF dynamics (black lines in Fig. 1) exhibited two (T = 10 – 120 s) or three (T = 180 – 300 s) local maxima. The highest ChlF maximum was always (except T = 10 s) appearing with the increasing light intensity and one may tentatively assume that this was reflecting a decrease of the photochemical quenching with the progressing congestion of the electron transport pathways in high light. This dominant maximum was followed by a drop in ChlF, likely due to an onset of the npq mechanisms. Since the position of the maximum was approximately coinciding with the light maximum for the periods T = 1 and 3 s, the photochemical quenching was decreased proportionally to light intensity, and npq was not fast enough to respond. With the period of oscillation T = 10 s, the main ChlF maximum appeared with a delay after the light maximum, indicating a mechanism other than a trivial decrease of photochemical quenching in homogeneous PSII. With the longer periods of the light oscillation (T = 20 – 300 s), the main maximum appeared sharper and peaking at or before the maximum light is achieved. One can tentatively presume onset of npq before the slowly increasing light reached its maximum. The smaller secondary maxima of ChlF were likely due to the cyclic electron transport (CET) or regulatory nonlinearity (see Introduction). To support our claims on the role of npq and CET, Fig. S4 in the supplementary material (data from Niu et al. 2022) show the forced oscillations in ChlF with period T = 60 s measured with Arabidopsis npq4 mutant (lacks PsbS-dependent npq; Li et al., 2000), and pgrl1ab mutant (lacks the main, antimycin-A sensitive, PGR5/PGRL1-dependent CET pathway; DalCorso et al., 2008). However, one can also consider that PSII as well as plastoquinone pool are heterogeneous (surveyed in Lazár, 2003). Thus, different kinetics of reduction of heterogeneous PSII and/or heterogeneous plastoquinone pool can contribute to appearance of more than one maximum in ChlF for periods of 10 s and longer.

The signal I830 (blue line) that is dominantly attributed to the oxidation of the P700 primary donor in PSI was dominantly following the light oscillation with two interesting additional dynamic features: distinct two dynamic peaks appearing around the period T = 20 s and a shift of the peaks to shorter times together with a decrease of the first peak with increasing period, the later leading with the periods T = 120 – 300 s to appearance of a minor, narrow secondary maximum that occurred in the low-light, in the early rising phase. These non-linear dynamic features may reflect P700 oxidation due to transient imbalance on the donor and acceptor sides of PSI. Considering that the I830 signal is to some extent affected by the redox changes in plastocyanin, and in a minor way also by ferredoxin, the observed secondary dynamic patterns might be related to a redox component other than P700. This is addressed in our other work (Niu et al., 2022) by using a new Walz instrument, which enables resolution of P700, plastocyanin, and ferredoxin. It should also not be ignored that PSI can be heterogeneous, with dynamically distinct pools in stroma lamellae and at the edges of the grana (e.g., Albertsson, 2001). The dynamic heterogeneity may also be expressed in distinct patterns during the oscillations.

None of the explanations that were tentatively proposed above brings a detailed insight into the dynamic phenomena and regulation that may be occurring in oscillating light and, thus, a more focused study is required. We choose for such a research the period T = 60 s, which is known to be characteristic for spontaneous oscillations in plants (e.g., Ferimazova et al., 2002; Lazár et al., 2005) and which was identified as a strong resonance period of a regulatory feedback in Nedbal and Březina (2002). First, the signals in panel T = 60 s of Fig. 1 were translated and shown in the input-output graphical presentation in Fig. 2, connecting to the earlier data in Nedbal et al. (2005). Clearly, all the three measured (output) signals respond non-linearly to the input signal (incident light intensity) and show a hysteresis: the signal dynamics in the ascending light phase differs from the descending light phase. The hysteresis is often explained in physics by existence of a memory effect, i.e. present state is influenced by the system history. With respect to kinetics in a simple reversible reaction, presence of a memory effect reflects the fact that backward reaction is slower than related forward reaction. For example, in the so-called mnemonical (or hysteretic) enzymes, their activation is much slower than their inactivation (Roussel, 1998), or, in photoprotection of photosynthesis by the npq, epoxidation of zeaxanthin to violaxanthin is slower than deepoxidation of violaxanthin to zeaxanthin (Matuszyńska et al., 2016). In a more complex system of reactions, the memory effect reflects an interplay of all involved reactions, including feed-forward and feed-back reactions. The strongest hysteresis is for P515, which confirms that it is a very integrative signal depending strongly on the past processes. A minimal hysteresis is observed in I830, reflecting the primary donor of PSI, which is largely light-driven by the primary photochemistry in the RC of PSI. ChlF signal exhibits a complex behaviour because it is a composite of primary photochemistry, redox state of, mostly, downstream electron transport chain, and the regulatory npq. Plastoquinone redox state is also influenced by CET that is inherently non-linear.

**Figure 2.**
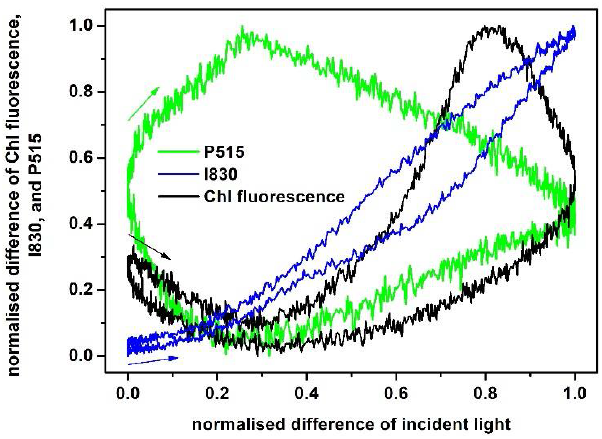
Dependence of normalized difference of chlorophyll (Chl) fluorescence, I830 and P515 signals on normalize difference of incident light intensity (input-output graph). The graph was constructed from data of Fig. 1 for the period T = 60 s. Position and direction of the colour arrows indicate way of the signals changes in the beginning of the light intensity period.

To separate individual factors contributing to the dynamics in Fig. 2, we used MTSPs of light that transiently congested the electron transport pathways and allowed quantitation of quantum yields in both PSs (Fig. 3). The changes in the effective quantum yield of PSI photochemistry (Y(I) in Fig. 3A) were antiparallel to the changes in the incident light intensity, reflecting oxidation of P700 by the strong light that reduced the yield. This process largely formed the dominant large peak in the I830 signal. The quantum yield of quenching of PSI excitation energy due to the limitation on the acceptor side of PSI (Y(NA) in Fig. 3A) was strongly modulated in the ascending low-light phase, while it seems to be hardly changing in other phases of the oscillation. The limitation on the PSI acceptor side is the reason why the effective quantum yield of PSI photochemistry is lower than unity in the beginning and in the end of the light period, and consequently that not all RC of PSI are open (see below) in the beginning and in the end of the light period. The quantum yield of quenching of the PSI excitation energy due to the limitation on the PSI donor side (Y(ND) in Fig. 3A) also contributes to the modulation of the I830 signal around the minor peak. Y(ND) has a small local maximum at the minor peak position, while Y(NA) has a local depression in the same phase, both probably signalling transiently reduced flow of electrons from the donor side of PSI. A largely dominant modulation of Y(ND) comes however later, close to the light maximum. Changes Y(ND) (Fig. 3A) follow the changes in the measured I830 signal. This is expected since the quenching is realized, when the donor side of PSI is oxidized and the I830 signals mostly reflects amount of P700^+^. We recall here the above-mentioned fact that the data presented in Fig. 3A might be distorted by the redox components other than P700.

**Figure 3.**
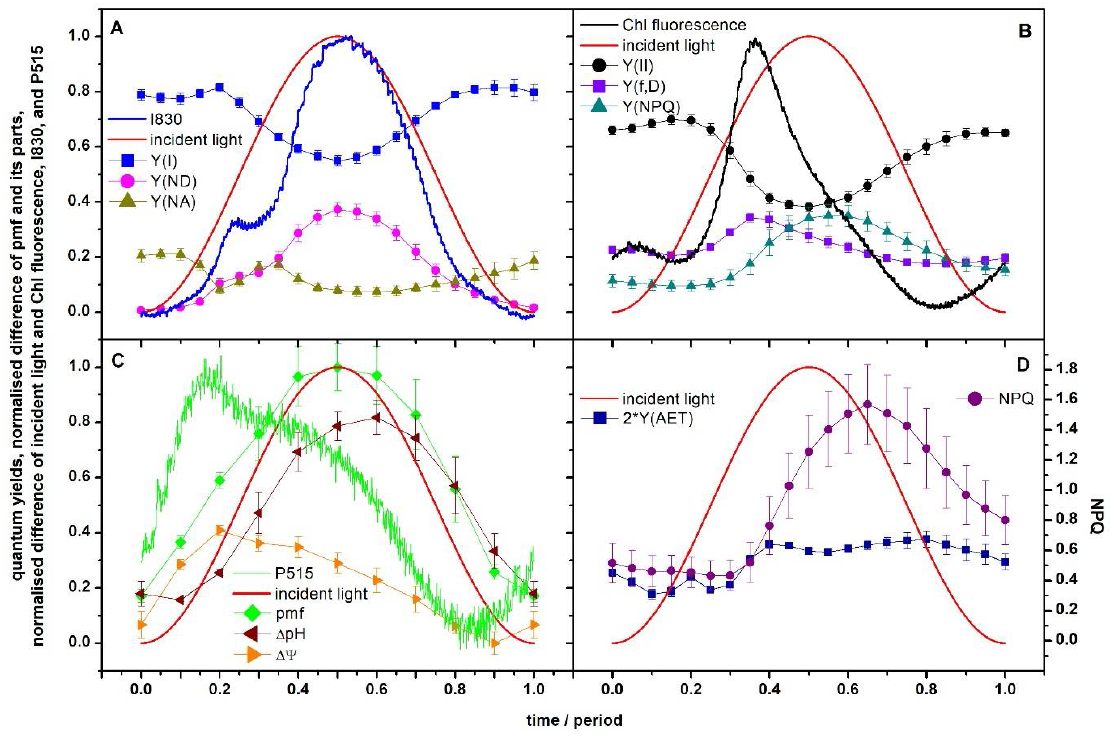
Parameters of photosynthetic energy partitioning during the forced oscillations with period of 60 s based on chlorophyll (Chl) fluorescence, I830 and P515 signals. (A) – effective quantum yield of PSI photochemistry (Y(I)) and quantum yields of non-photochemical quenching of PSI excitation energy due to limitation at the PSI donor (Y(ND)) and acceptor (Y(NA)) sides, (B) – effective quantum yield of PSII photochemistry (Y(II)) and quantum yields of constitutive non-regulatory (Y(f,D)) and of light-induced regulatory (Y(NPQ)) non-photochemical quenching of PSII excitation energy, (C) – relative pmf, ΔpH-dependent part of pmf and ΔΨ (range of values of pmf and its two parts are normalized to 0-to-1 range), and (D) – effective quantum yield of alternative electron transport (Y(AET)) multiplied by two (for a better visualisation) and the NPQ parameter (right y-axis). Courses of normalized difference of the incident light and of the signals are also shown. The symbols represent mean values (n = 3 – 4) and the error bars (sometimes hidden by the symbols) show standard deviations. For more details, see Material and methods.

The quantum yields related to function of PSII are shown in Fig. 3B; these data are based on the measured ChlF values. An alternative evaluation based on a correction of F_M_ and F_M_’ due to a contribution of variable ChlF originating from the closed RCIIs (see Introduction) is presented in the supplementary material in Fig. S5. The effective quantum yield of PSII photochemistry Y(II) (Fig. 3B) displayed, similarly to Y(I) (Fig. 3A), a pattern that was roughly antiparallel to the light oscillation, only slightly delayed in phase. The delay indicated memory effects in the PSII photochemistry. Memory was likely causing also the pattern asymmetry (hysteresis); the decline of Y(II) when the light was increasing was steeper than the increase when the light is decreasing in the second half of the light period.

The quantum yield of regulatory npq of the PSII excitation energy (Y(NPQ) in Fig. 3B) followed with a delay the light oscillation, forming a wide maximum in high light around mid-period that persisted long into the descending light phase. The quantum yield of the constitutive non-regulatory npq of PSII excitation energy (Y(f,D)) in Fig. 3B) was similar to the dynamics of ChlF (Fig. 3B). This is expected, since the rate constants of the constitutive non-regulatory non-photochemical quenching and of the ChlF are similar and in mathematical models often joined together into one value (Lazár, 2015) and thus changes in Y(f,D) and ChlF should follow the same trend. Similarly to PSI (Fig. 3A), the constitutive (Y(f,D)) and the regulatory (Y(NPQ)) npq are the reasons, why the effective quantum yield of PSII photochemistry (Y(II)) is lower than unity in the beginning and in the end of the light period.

The values of Y(II) stay lower than the values of Y(I) throughout the entire illumination period. A difference between Y(I) and Y(II) is ascribed to the alternative electron transport (AET) (e.g., Yamori et al., 2011). By the AET we mean all electron transport pathways except the linear one (PSII → plastoquinone pool → cytochrome b_6_/f → plastocyanin → PSI → ferredoxin → NADP^+^). The effective quantum yield of AET (Y(AET)) multiplied by two (to better visualize the changes) is shown in Fig. 3D (left y-axis). The Y(AET) varies between ca. 0.1 and 0.2, with the lower values in the first half period when the light is rising. The variation of Y(AET) reflects an interplay of multiple pathways (reviewed in Alric and Johnson, 2017).

Figure 3D also shows the npq parameter (NPQ; right y-axis). Since NPQ = Y(NPQ)/Y(f,D), it represents a ratio between the rate constant for light-induced regulatory npq of PSII excitation energy and the rate constant of constitutive non-regulatory npq of the PSII excitation energy. It is evident that the rise and decrease of the NPQ parameter is delayed with respect to changes in the incident light intensity. This delay reflects time constant of activation of the regulatory npq of PSII excitation energy, which takes in our case about 15% of the period, i.e., about 9 seconds. Further, the changes in NPQ parameter roughly occur at about the same oscillation phase as that of Y(AET). This can be understood by considering that AET includes as a component also the CET around PSI (reviewed in Alric and Johnson 2017) and that CET acidifies the lumen, a prerequisite for the npq regulation (reviewed in Holzwarth and Jahns, 2014). Parallel dynamics of Y(AET) and NPQ parameter are, thus, conceivable.

The relative changes of the pmf and of its ΔpH-dependent and ΔΨ components (not as fractions of pmf, see Material and methods for details) during the forced oscillations were obtained by analysing the P515 signal. The oscillating illumination was changing pmf as well as the contributions of its chemical and electrical components and all three are shown in Fig. 3C together with the P515 signal. The pmf followed approximately the light intensity, with apparent deviations at the beginning and the end of the light period. The ΔpH-dependent part of pmf was delayed relative to light by ca 1/10 of the period (≈ 6s). With ΔpH being driven by the water splitting in PSII, oxidation of plastoquinol by cytochrome (cyt) b_6_/f, and, in opposite direction, by the CF_0_-CF_1_ ATP-synthase, the observed delay of 6 s may reflect an interplay of all these processes. The NPQ parameter (Fig. 3D) is modulated by the oscillating light in a pattern that is similar but phase-shifted to the ΔpH-dependent part of pmf (Fig. 3C). The phase delay confirms that the lumen acidification (reflected in ΔpH) is not the only driver of the light-induced npq (reviewed in Holzwarth and Jahns, 2014).

The dynamic patterns of ΔΨ and P515 are similar, which is not surprising since the P515 signal is a measure of the electrochromic shift of pigments due to the ΔΨ (see Introduction). The ΔΨ initially increases with increasing incident light intensity with a maximum of ΔΨ at 0.2 of the period. From this maximum, ΔΨ then decreases even though the incident light intensity continuous increasing. We tentatively propose the flux of other positive ions (K^+^, Mg^2+^) from lumen to stroma and of negative ions (Cl^-^) from stroma to lumen, that all counteract charge of accumulated protons in lumen (see, e.g., Cruz et al., 2001; Davis et al., 2017; Lyu and Lazár, 2017; Li et al. 2021) are responsible for this dynamic feature in ΔΨ.

For further insight into the dynamics of the regulation, we also determined from ChlF the coefficient of photochemical quenching of PSII excitation energy, qCU (Fig. 4), which reflects fraction of open PSII RCs (RCIIo) in the light-acclimated state, assuming the energetic connectivity among PSII units (Kramer et al., 2004, reviewed in Lazár, 2015). In the first quarter period when the light already rises strongly, qCU changes only a little. It starts dropping, PSII RCs closing sharply, only shortly before the maximum light is reached. The subsequent reopening of the PSII RC occurs gradually with decreasing light in the second half period (Fig. 4). We note that fraction of RCIIo can be estimated also by means of qP or qL (see Kramer et al. 2004, Lazár 2015), which are estimates for RCIIos within the separated units model (given PSII and its antenna are energetically separated from other PSIIs and their antennas) or the lake model (particular PSIIs share excitations from all antennas without any restrictions), respectively. The qP and qL courses show similar pattern to qCU but the values are different reflecting the assumed energetic communication among the PSII units (Fig. S6 in the supplementary material).

**Figure 4.**
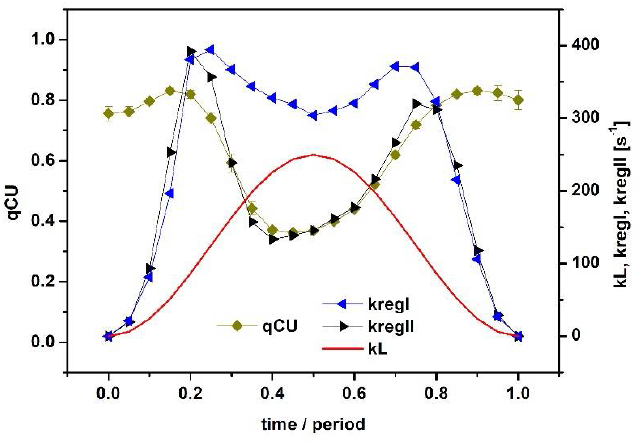
Courses of coefficient of photochemical quenching of PSII excitation energy, qCU, and of the rate constants, kL, kregI, kregII. The qCU reflects fraction of open PSII RCs, kL reflects rate constant of incident light intensity, and kregI and kregII reflect apparent rate constants of all regulatory mechanisms causing re-opening of PSI and PSII, respectively, during the forced oscillations with period of 60 s. The symbols for qCU represent mean values (n = 4) and their error bars (standard deviations) are sometimes hidden by the symbols. For more details, see the text.

We further wanted to quantify at which phases of the light period a regulation occurs. For that purpose, we assumed that the RCIIos are converted by light to closed PSII RCs (RCIIc) with a rate constant kL that is proportional to the incident light intensity. The action of npq, which in fact lowers kL, as well as of all other regulatory mechanisms causing apparent re-opening of RCIIcs, is described by an apparent rate constant of PSII regulation, kregII. In the framework of this simple evaluation, it is the same if the regulation causes decrease of kL or is considered as an increase of kregII. Since kL can be estimated from the known incident light intensity (see Material and methods), we can estimate kregII as follows:

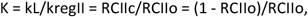

and then, by approximating RCIIo by qCU we can write:

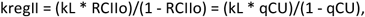

where K is an equilibrium constant. This approach is based on an assumption that the system is in a steady state equilibrium at every time of measurement. The assumption is fulfilled in our experiments since the forced oscillations were stable with a sustained pattern.

Similarly with PSI. According to the theory of PSI quantum yields (Schreiber and Klughammer, 2008a), RC of PSI is open (RCIo), when the donor side of PSI is reduced and at the same time the acceptor side of PSI is oxidised. Consequently, fraction of RCIo numerically equals to Y(I) (Fig. 3A). Assuming that kL is the same for PSII and PSI, we can write for the apparent rate constant of PSI regulation, kregI:

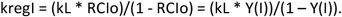

Changes of the regulation rate constants, kregII and kregI during the forced oscillations with period of 60 s are shown in Fig. 4. Both kregII and kregI have two peaks at the same position; at about 0.2 and of the period, close to the inflection points of the light modulation. The regulation buffers the variations of RCIIo (qCU in Fig. 4) and of RCIo (Y(I) in Fig. 3A) in the initial ascending and final descending phases of the oscillating incident light intensity. Thus, the regulations try to keep both RCIIs and RCIs open although the light change is fastest at the inflection points. Only when the further light increase towards the maximum exceeds the regulation capacity, the RC’s close (dips in qCU in Fig. 4 and in Y(I) in Fig. 3A at about 0.5 of the period). We note that an alternative evaluation of kregII using qP or qL (see above) for estimation of fraction of RCIIo leads to the same qualitative results; the positions of the regulation maxima are the same as based on qCU but the values of kregII are lower for qL or higher for qP (Fig. S7 in the supplementary material).

## Discussion

In contrast to the spontaneous oscillations studied extensively in the eighties of the previous century (reviewed in Walker, 1992; Giersch, 1994), studies reporting on the forced oscillations remain rare. Aiming at exploiting full potential of forced oscillations, the conceptual background was recently newly formulated in Nedbal and Lazár (2021). We note that to the spontaneous oscillations occur only under extreme conditions; either high CO_2_ concentration or strong actinic light must be applied. On the other hand, the forced oscillations can be measured in ambient CO_2_ concentration and in a range of light intensities, which occur in the field. Thus, the forced oscillations reflect function and regulation of photosynthesis under natural conditions.

The results reported here complement the previous studies that mostly focused at ChlF and related parameters: Y(PSII), Y(f,D), Y(NPQ), NPQ, F_V_’/F_M_’ (Nedbal and Březina, 2002; Nedbal et al., 2003; 2005; Nedbal and Lazár, 2021; Shimakawa and Miyake, 2018; Samson et al., 2019), at CO_2_ assimilation and I830 signal (Nedbal et al., 2003; Shimakawa and Miyake, 2018), and at I830-related parameters: Y(PSI), Y(ND), Y(NA) (Shimakawa and Miyake, 2018). The novelty here is in reporting on the forced oscillations in the P515 signal (green lines in Figs. 1 and 2) and on the relative changes of the pmf and its ΔpH-dependent part and ΔΨ (Fig. 3C), both determined from the P515 signal. Further, rate constants of regulation of open RCIIs and RCIs, kregII and kregI, respectively, are calculated as they change in oscillating light (Fig. 4). This analysis shows that regulation acts to stabilize the fraction of open RCs even though the light oscillates. In other words, the photosynthetic regulation acts towards stable output with fluctuating input, i.e., stable fraction of RCIIo and RCIo in oscillating light. This conclusion was tentatively proposed earlier (Nedbal et al., 2005) based on the input-output relation of the ChlF on oscillating light and is confirmed here by solid experimental evidence. Similar conclusion has been recently revealed behind the absorption spectra of photosynthetic pigments (Arp et al., 2020).

The npq would be the most common candidate to explain the regulation of PSII. This is supported by a difference between ChlF of wild-type and the npq4 Arabidopsis mutant (Fig. S4; Niu et al., 2022), which lacks PsbS-dependent npq (Li et al., 2000), the differences being most pronounced in the second half of the period. This in an agreement with the fact that the NPQ parameter has a high value (Fig. 3D), but not a maximum, at position of the second maximum of kregII (Fig. 4) but the NPQ parameter has a low value at position of the first maximum of kregII. Thus, the npq alone, initiated by accumulation of protons in the lumen reflected in increase of the ΔpH-dependent component of the pmf (Fig. 3C), cannot explain all changes in kregII. On the other hand, position of the first maximum of kregII and kregI (Fig. 4) is the same as position of the maximum of ΔΨ (Fig. 3C). ΔΨ has been reported to promote charge recombinations in PSII (e.g., Dau and Sauer, 1992; Davis et al., 2016), causing thus opening of RCIIs. If the recombinations are nonradiative (Davis et al., 2016), they would lead to decrease of ChlF. In fact, the ChlF slightly decreases (Fig. 3B) at position of the first maximum of kregII (Fig. 4). Thus, we suggest role of ΔΨ in regulation at least of PSII in the rising phase of sinusoidal illumination. A role of ion fluxes in fluctuating light has been reported previously (e.g., Duan et al., 2016; Li et al., 2021). The above mentioned roles of the ΔΨ and ΔpH-dependent components of pmf on PSII regulation in different parts of the oscillating light period are in agreement with the role of the pmf partitioning in the regulation (Avenson, et al. 2004; Takizawa et al., 2007; Davis et al., 2017).

The CET from PSI back to the PQ pool has two pathways: sensitive and insensitive to antimycin A. The PGR5/PGRL1 complex (e.g., Munekage et al., 2002) and the NAD(P)H-dehydrogenase-like complex (e.g., Yamamoto et al., 2011) are involved in the sensitive and insensitive pathways, respectively. The CET is often considered as a regulatory mechanism for electron transport in the TM (reviewed in Alric and Johnson, 2017) and its protective role, especially of the PGR5/PGRL1-dependend CET, under fluctuating light has also been reported (e.g., Yamori et al., 2016; Yamamoto and Shikanai, 2019). The CET promotes also the so-called photosynthetic control, i.e., a decrease of the rate of reduced plastoquinone oxidation at cyt b_6_/f due to accumulation of protons in the lumen (e.g., Johnson and Berry, 2021), the effect being also known as backpressure of protons (Siggel 1976). However, we can exclude CET as origin of the peaks in course of kregII and kregI, since the CET would partly re-open closed RCIs, compared to the case without any CET, but, by reduction of the PQ pool, also promoted by the photosynthetic control, the CET would contribute to increased closure of RCIIs, which is against simultaneous changes of RCIIo (qCU in Fig. 4) and of RCIo (Y(I) in Fig. 3A) in the same direction. As mentioned above, the regulation is maximal at low intensities of excitation light and, on the other hand, some AET pathways were inferred to be functional only at low light intensities, e.g., malate valve and Mehler reaction (Walker et al., 2014). Mehler reaction has been suggested to play an important role in photoprotection of PSI under fluctuating light (Sun et al., 2020). In our case, the electron flow from PSI to malate and/or oxygen would re-open PSI and consequently also PSII, which is in agreement with simultaneous changes of RCIIo (qCU in Fig. 4) and of RCIo (Y(I) in Fig. 3A) in the same direction and consequently of kregII and kregI. Thus, we tentatively assign the regulation to function of a part of AET. In fact, there is a small peak in Y(AET) at 0.2 of the period and a broad peak in Y(AET) at 0.8 of the period (Fig. 3D), i.e., at the same times as the maxima of kregII and kregI (Fig. 4). Thus, as mentioned above, a part of AET, the CET around PSI, contributes to regulation of the npq, and another part of AET, probably malate valve and/or Mehler reaction, contributes to regulation of RCs opening. In PSII, the effort keeping the RCIIs open might be additionally caused by the npq and the ΔΨ-promoted charge recombinations (see above).

The above discussion is on our data obtained by means of a standard measurement and evaluation. However, as mentioned in the Introduction, also RCIIc can emit variable ChlF, the effect playing a role when the saturating pulses are applied, which has been also our case. Thus, we have also performed an alternative evaluation, which employs a correction of ChlF values obtained upon saturating pulses (the F_M_ and F_M_’ values). The correction is described in the supplementary material and the results are shown in Figs S5, S8, S9 ibid. This alternative evaluation leads to changes in values of related parameters but their qualitative changes during the oscillating light period is the same as using the standard evaluation. Thus, this alternative evaluation does not change our conclusions. However, this alternative evaluation of the ChlF parameters, together with discrimination of the components (P700^+^, oxidised plastocyanin, reduced ferredoxin) contributing to the I820 signal, using DUAL-KLAS-NIR instrument by Walz, might bring new conclusions. This will be done in our future work.

Even if we have not used any mutant, which lacks or overexpresses particular protein(s) involved in the regulation of photosynthesis, by combined measurement of ChlF, I830, and P515 reporter signals, and evaluation of related parameters describing energy partitioning in PSI, PSII and of the pmf, we were able to infer the mechanisms of regulation of photosynthesis in fluctuating, in our case sinusoidal light. The forced oscillations measured with different mutants will be presented by us elsewhere (Niu et al., 2022), a part of this data being shown now in the supplementary material. Our work also shows a high potential of the forced oscillations in studying function and regulation of photosynthesis. However, the above discussion demonstrates complexity of the problem of photosynthetic regulation. Further progress in understanding the forced oscillations and the regulations is expected from experiments involving chemical interventions by electron acceptors and donors, as well as inhibitors and with mutants affected in well-defined regulatory mechanisms or pathways. This will stimulate falsification and further development of structure-function-based mathematical models considering particular regulatory mechanisms. In the opposite top-down direction, decoding the forced oscillations will also be supported by system identification and system control, tools that have already been successfully applied to solve homologous challenges in engineering (e.g., Schrangl et al., 2020) and medicine (e.g., Doyle, 2016).

## Abbreviations

Chl: chlorophylll
ChlF: chlorophyll fluorescence
ΔpH: difference of pH across TM
ΔΨ: difference of electric potential across TM, the membrane potential
npq: non-photochemical quenching
pmf: proton motive force
RC: reaction centre
TM: thylakoid membrane

## Acknowledgement

D.L. was supported from ERDF project “Plants as a tool for sustainable global development” (no. CZ.02.1.01/0.0/0.0/16_019/0000827). L.N. gratefully acknowledges the financial support by the Federal Ministry of Education and Research of Germany (BMBF) in the framework of YESPVNIGBEN project (03SF0576A).

## Author contributions

D.L. conceived the study, did the experiments, analysed the experimental data, and wrote the manuscript. L.N. prepared knowledge foundation for experimental design and wrote the manuscript. Y.N. measured the Arabidopsis data.

## Conflict of Interest

The authors declare no conflict of interest.

## Data availability

The data supporting the findings of this study are available from the corresponding author, (Dušan Lazár), upon request.

## Supplementary material

Data presented in Figs. S5, S8, and S9 represent an alternative evaluation of data presented in Fig. 3B of the main text and in Figs. S6 and S7 of the supplementary material, respectively. The alternative evaluation considers a correction of F_M_ and F_M_’ whose measured values might be overestimated by variable ChlF of closed reaction centres of PSII (Vredenberg and Bulychev, 2002; Magyar et al., 2018; Laisk and Oja, 2020; Sipka et al., 2021). Thus, instead of the measured F_M_ and F_M_’, corrected F_M_ and F_M_’ values were used. Data presented in Laisk and Oja (2020) show that about 1/3 of ChlF measured with multiple turnover saturating light pulse (F_M_ and F_M_’), as it is used in our measurements, is caused by increase of ChlF of already closed reaction centres of PSII. Thus, we considered 2/3 of the measured F_M_ and F_M_’ as the corrected values of F_M_ and F_M_’. These corrected values were then used for evaluation of the quantum yields of function of PSII (Fig. S5) and of the coefficients of the photochemical quenching of ChlF, qP, qCU, and qL (Fig. S8). Definition of qCU s provided in the main text and qP and qL were calculated as follows (Kramer et al., 2004; Lazár, 2015):

- qP = (F_M_’ - Ft)/(F_M_’ - F_0_’)
- qL = qP * (F_0_’ / Ft),

where the ChlF values have the meaning defined in the main text.

**Figure S1.**
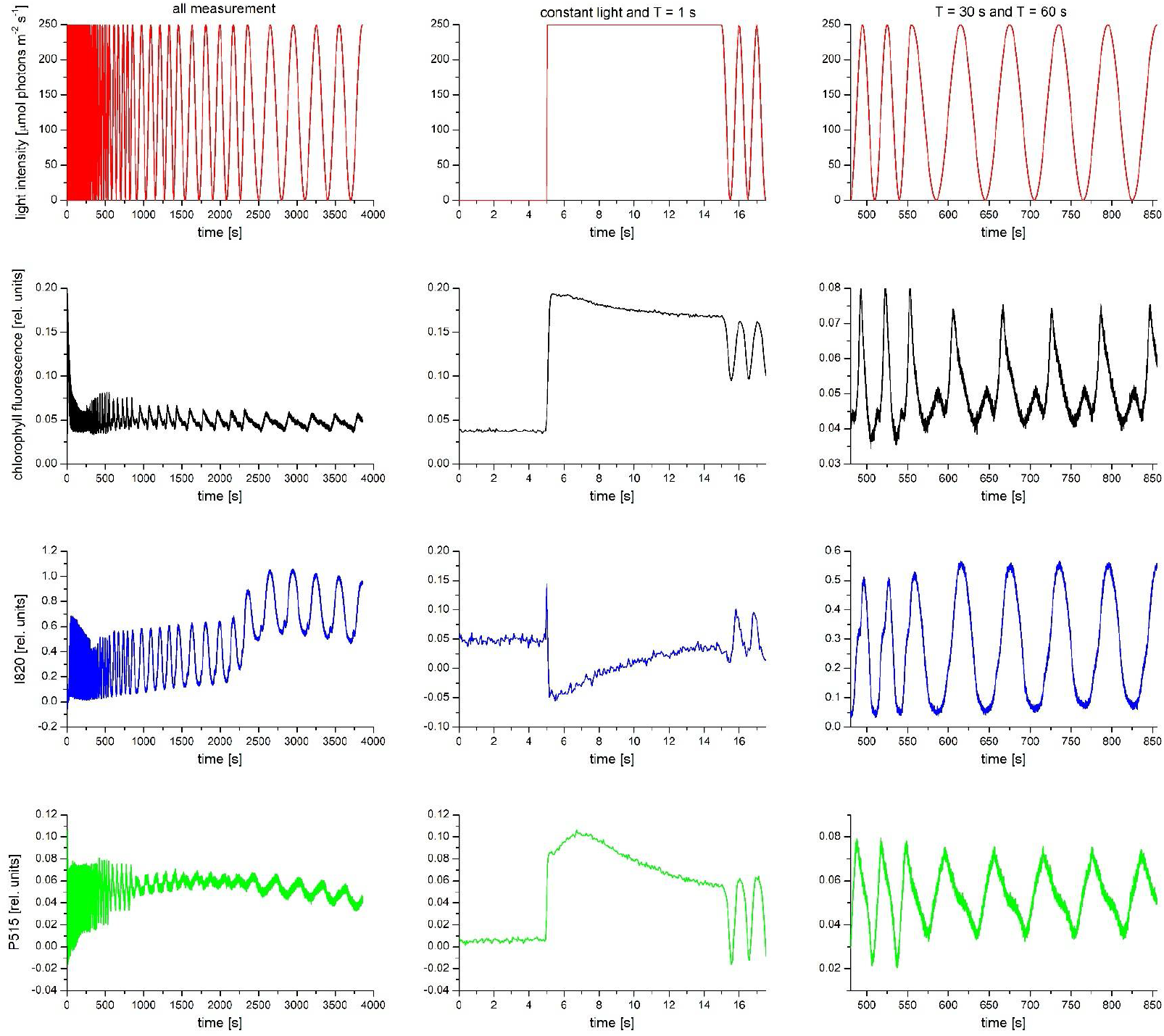
Example of raw data of typical measurements of oscillations of ChlF (second row), I820 (third row), and P515 (fourth tow) forced by sinusoidally oscillating incident light (first row). The first column shows the whole measurement; it mainly serves to show a drift of the baselines in the I820 a P515 signals (see Material and methods). The second and third columns show in detail selected time intervals of the whole measurement. It shows how the constant light was changed to the oscillating light with period T of 1 s (two and half 1-s-periods are shown; second column), and last two and half periods of 30 s changed to five periods of 60 s (third column). The data also show that shape of particular oscillations for given period of forcing has not changed during time. Four such whole measurements have been done, each with different leaf from different plant. The data presented in the Fig. 1 in the main text have been obtained by averaging two last whole periods.

**Figure S2.**
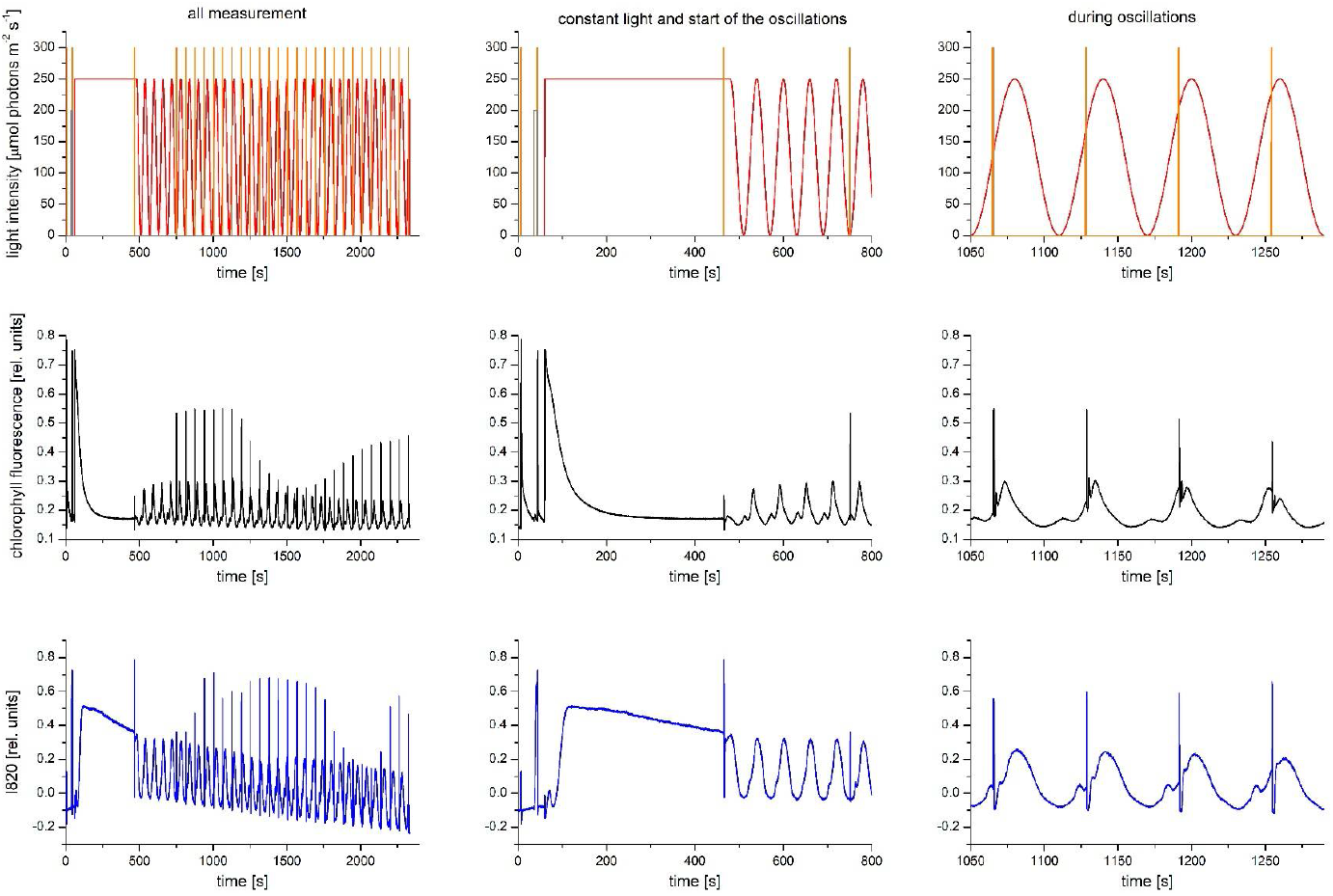
Example of raw data of typical measurements of quantum yields of PSII and PSI during oscillations of ChlF (second row) and I820 (third row) signal, respectively, forced by sinusoidally oscillating incident light (first row) with period T = 60 s. The grey and orange vertical lines show positions of the far-red light illumination (necessary to determine P_M_) and of the MTSPs, respectively (see Material and methods), and the heights of the lines are not in scale with respect to intensity of the oscillating light. The first column shows the whole measurement; it serves to show a small drift of the baseline in the I820 signal, which does not affect evaluation of the quantum yields, and to show magnitudes of the ChlF and I820 signal caused by the MTSPs compared to the magnitude of the oscillation itself. The second and third columns show in detail selected time intervals of the whole measurement. Namely, the second column shows the first five and half oscillation periods after the constant light illumination; it demonstrates that three periods were enough to obtain the same shape of the oscillations over time. The third column shows shift of the positions of the MTSPs by 3 s at each oscillation period. Thus, (T/3) + 1 = (60/3) + 1 = 21 oscillation periods/MTSPs were necessary to measure the yields with 3-s-step during the whole 60-s-period. The same shape of the oscillations over whole measurement show that one MTSP applied during one oscillation period had no effect on the shape of following oscillations. Three such whole measurements have been done, each with different leaf from different plant. The data presented in the Fig. 3A,B in the main text were obtained by merging responses to the shifted MTSPs in 21 oscillation periods to one oscillation period.

**Figure S3.**
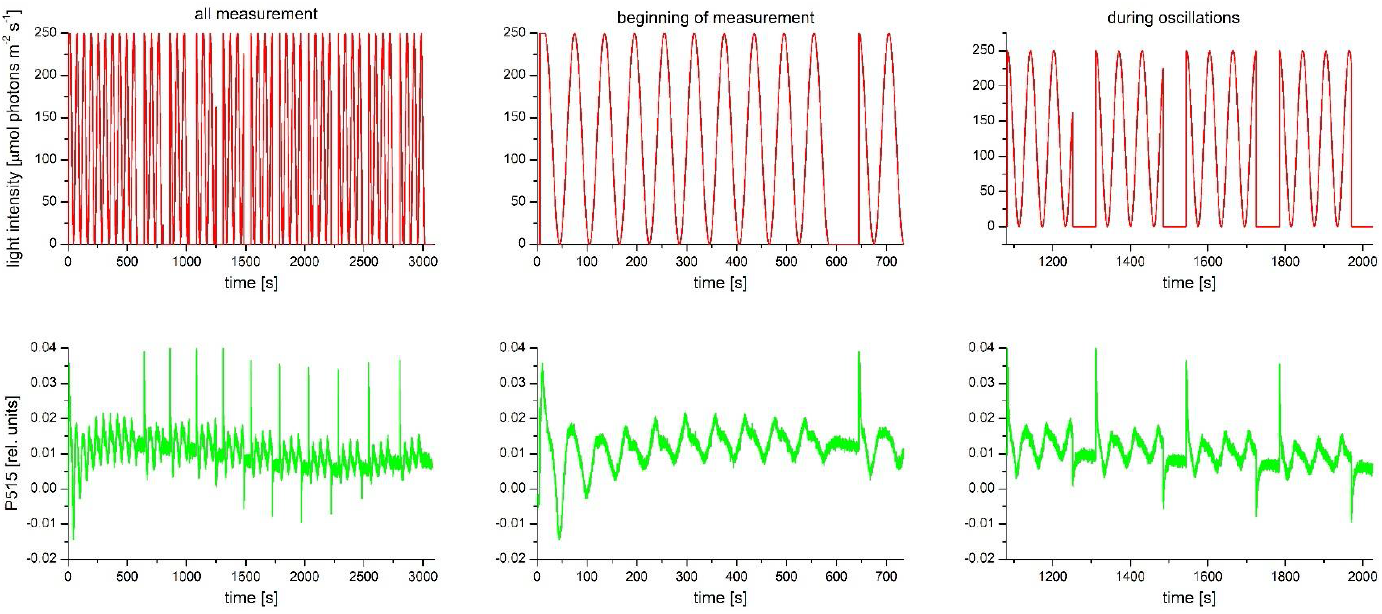
Example of raw data of typical measurement of the pmf partitioning into its ΔpH- and ΔΨ-dependent parts during oscillations of P515 signal (second row) forced by sinusoidally oscillating incident light (first row) with period T = 60 s. The partitioning was determined by switching off the light for 60 s at different phases of the oscillating light (for details, see Material and methods). The first column shows the whole measurement; it serves to show a small drift of the baseline in the P515 signal, which does not affect evaluation of the partitioning. The second and third columns show in detail selected time intervals of the whole measurement. Namely, the second column shows the first nine and half oscillation periods after 10-s constant light illumination; it demonstrates dynamics to reaching a stationary oscillatory pattern in P515 signal. The first 60-s dark interval, which started just in the beginning of the 60-s oscillating light, followed by one and half 60-s light periods are also shown. The third column shows four 60-s dark intervals each subsequently shifted 6 s forward in the oscillating light period (for details, see Material and methods). Thus, (T/6) + 1 = (60/6) + 1 = 11 60-s dark intervals were necessary to measure the partitioning with 6-s-step during the whole 60-s-period. The same shape of the oscillations before beginning of the 60-s dark interval over whole measurement show that two and half light periods following the 60-s dark interval were enough to obtain stationary oscillatory pattern. Three such whole measurements have been done, each with different leaf from different plant. The data presented in the Fig. 3C in the main text have been obtained by merging responses to the shifted 60-s dark interval in 11 oscillation periods to one oscillation period.

**Figure S4.**
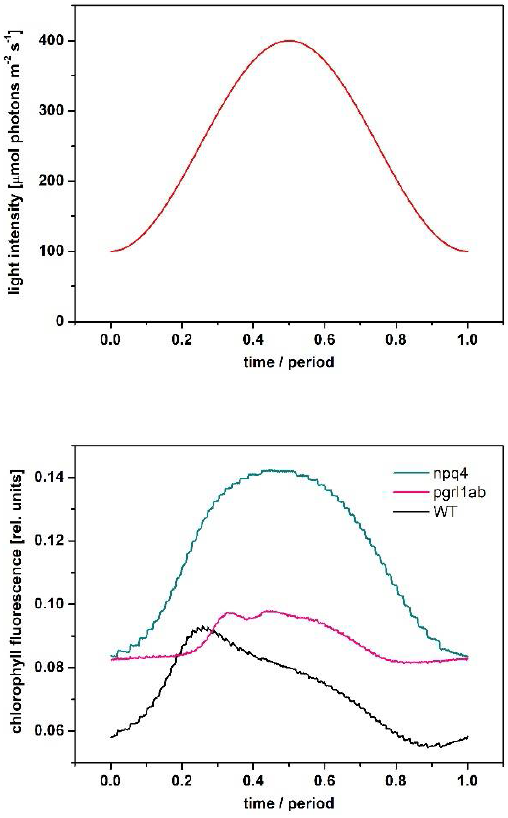
Example of raw data of typical measurements of ChlF forced by sinusoidally oscillating incident light with period T = 60 s. The measurements have been done with wild type and npq4 (lacks PsbS-dependent npq; Li et al., 2000) and pgrl1ab (lacks the main, antimycin-A sensitive, PGR5/PGRL1-dependent CET pathway; DalCorso et al., 2008) Arabidopsis mutants (data from Niu et al. 2022) for light intensity oscillating between 100 and 400 mmol photons m^-2^ s^-1^. The shape of the ChlF oscillation of the wild type is qualitatively similar to the shape of the oscillation with pea for the same period. The data show that ChlF of the npq4 mutant almost copies the shape of oscillation of the light intensity, suggesting a role of the npq in decrease of ChlF after its maximum observed in the wild type and in pea. On the other hand, ChlF of the pgrl1ab mutant is unchanged in the first approx. 12 seconds when the light intensity is quickly growing, suggesting a role of the CET in the initial increase of ChlF in wild type and in the small secondary maximum of ChlF in pea.

**Figure S5.**
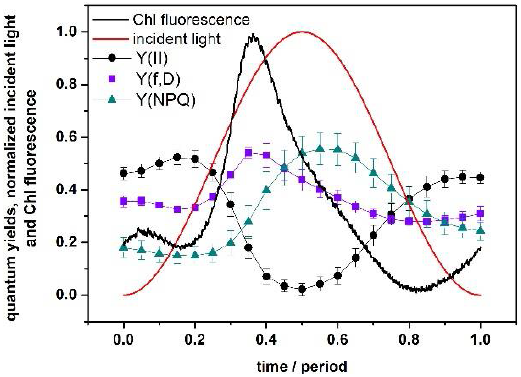
Parameters of photosynthetic energy partitioning during the forced oscillations with period of 60 s based on ChlF, namely effective quantum yield of PSII photochemistry (Y(II)) and quantum yields of constitutive non-regulatory (Y(f,D)) and of light-induced regulatory (Y(NPQ)) non-photochemical quenching of PSII excitation energy. The quantum yields were evaluated considering a correction of the F_M_ and F_M_’ values (see beginning of the supplementary material), whose measured values might be overestimated by variable ChlF of closed reaction centres of PSII. Courses of the normalized incident light and of ChlF are also shown. The symbols represent mean values (n = 4) and the error bars (sometimes hidden by the symbols) show standard deviations.

**Figure S6.**
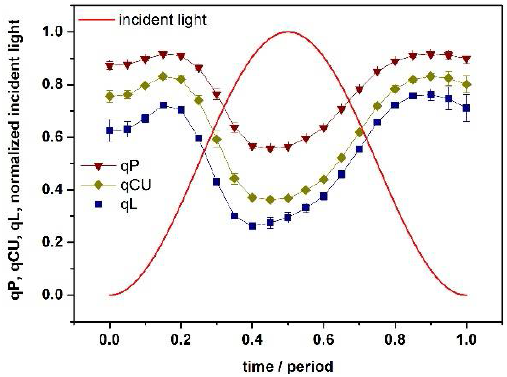
Courses of coefficients of photochemical quenching of excitation energy of PSII, qP, qCU, and qL, during the forced oscillations with period of 60 s. Course of normalized incident light intensity is also shown. All the coefficients can be used for estimation of fraction of the open reaction centres of PSII but assuming different energetic “communication” among PSII units; the units are energetically separated (qP), are energetically connected with a restriction (qCU), or are energetically connected without any restriction (qL). The symbols represent mean values (n = 4) and the error bars (sometimes hidden by the symbols) show standard deviations.

**Figure S7.**
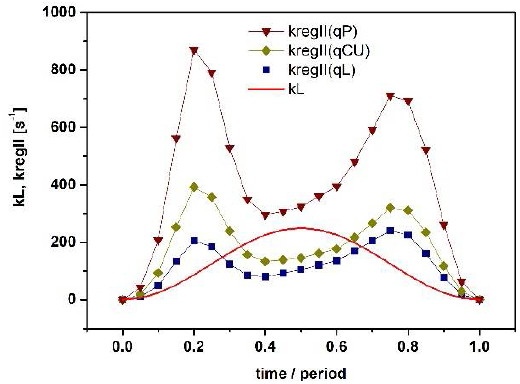
Courses of rate constants kL and kregII during the forced oscillations with period of 60 s. The kL reflects incident light intensity and kregII reflects apparent rate constant of all regulatory mechanisms causing re-opening of photosystem II reaction centres. Values of kregII were calculated from courses of qP, qCU, and qL shown in Fig. S6.

**Figure S8.**
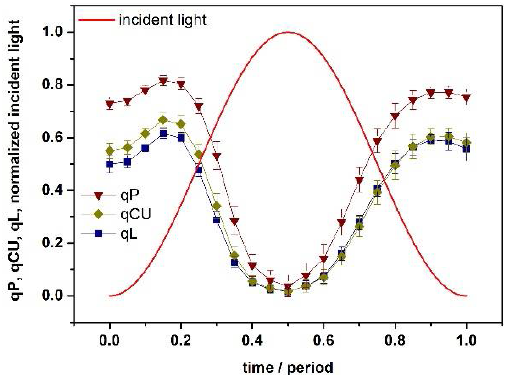
Courses of coefficients of photochemical quenching of excitation energy of PSII, qP, qCU, and qL, during the forced oscillations with period of 60 s. Course of normalized incident light intensity is also shown. All the coefficients can be used for estimation of fraction of the open reaction centres of photosystem II but assuming different energetic “communication” among PSII units; the units are energetically separated (qP), are energetically connected with a restriction (qCU), or are energetically connected without any restriction (qL). The coefficients were evaluated considering a correction of the F_M_ and F_M_’ values (see beginning of the supplementary material), whose measured values might be overestimated by variable ChlF of closed reaction centres of PSII. The symbols represent mean values (n = 4) and the error bars (sometimes hidden by the symbols) show standard deviations.

**Figure S9.**
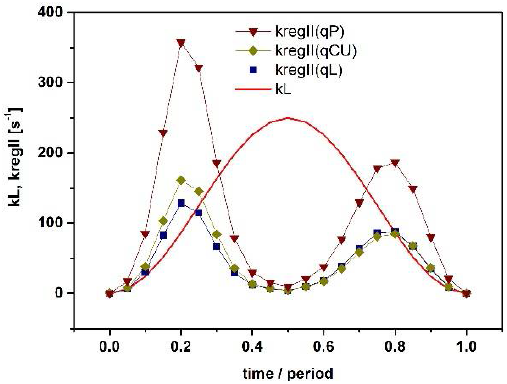
Courses of rate constants kL and kregII during the forced oscillations with period of 60 s. The kL reflects incident light intensity and kregII reflects apparent rate constant of all regulatory mechanisms causing re-opening of PSII reaction centres. Values of kregII were calculated from courses of qP, qCU, and qL shown in Fig. S8.

## Notes

### Competing Interest Statement

The authors have declared no competing interest.

